# L-Arginine Hydrochloride Enables Cold-Chain-Free DNA Polymerase Storage

**DOI:** 10.1101/2025.05.16.654417

**Authors:** Joseph Shenekji, Antonius Al-Daoud, Abduljalil Ghrewati

## Abstract

Preserving DNA polymerases under ambient conditions is a critical challenge, particularly in resource-limited settings with minimal cold storage infrastructure. This study introduces a novel, cost-effective approach to polymerase stabilization using L-arginine hydrochloride (LAH), evaluating its efficacy for Pfu polymerase and polymerase mixtures stored in 20 µl aliquots at 4°C, 25°C, and 37°C over three months. Enzyme activity was assessed via polymerase chain reaction (PCR) amplification of a 1250 base pair (bp) DNA fragment. Pfu polymerase maintained enzymatic activity at all tested temperatures with 1 M LAH, whereas activity was lost at 37°C without the stabilizer. LAH also enhanced the stability of Taq polymerase and Pfu + Taq mixtures, though with reduced efficacy. Mechanistically, LAH prevents protein aggregation, enhances solubility, and stabilizes enzyme structure through electrostatic interactions. These findings position LAH as a robust stabilizer, reducing cold chain dependency and enabling scalable molecular diagnostics in low-resource environments. This study advances enzyme preservation by offering a sustainable alternative to traditional methods, with applications in clinical diagnostics and biotechnology. Future work will optimize LAH concentrations, assess long-term stability beyond three months, and extend the approach to other thermostable polymerases.

## 1. Introduction

L-arginine, a naturally occurring amino acid, has emerged as a promising agent for preserving polymerase enzymes at ambient temperatures, addressing the critical need for stable biochemical reagents in research, diagnostics, and industrial applications [13]. The preservation of enzyme activity under ambient conditions has been a long-standing challenge in biotechnology, with early studies highlighting the need for stabilizers to prevent thermal inactivation [11]. L-arginine’s cationic properties not only inhibit microbial growth but also significantly enhance the stability and functional longevity of polymerases, which are essential for nucleic acid amplification and other biochemical processes [15]. L-arginine derivatives, such as L-arginine hydrochloride (LAH) and arginine ethyl ester (LAE), are widely utilized in the food and pharmaceutical industries to maintain enzyme activity and ensure product safety [20]. As a naturally occurring compound in living organisms, LAH offers a safer alternative to synthetic preservatives, which are often associated with health and environmental concerns, providing effective antimicrobial action without adverse effects [5]. The flexibility of LAH usage concentrations allows its application in diverse contexts, from food preservation to enzyme storage solutions [4].

Research has demonstrated that LAH significantly improves the storage stability of polymerase enzymes, enabling effective preservation at ambient temperatures and reducing dependence on energy-intensive low-temperature storage systems [2]. This capability is particularly valuable for low- and middle-income countries, where cold storage infrastructure is often limited, facilitating equitable access to molecular reagents and diagnostics during medical emergencies, such as the COVID-19 pandemic, which highlighted logistical challenges in PCR reagent and vaccine distribution [8, 19, 22]. These challenges include logistical constraints requiring specialized equipment (e.g., -20°C freezers, refrigerators, and cold packs), transportation difficulties in remote areas, and high operational costs, as outlined by global health guidelines [22]. Additionally, ambient temperature storage reduces the environmental impact of refrigeration and cold-chain freight, aligning with sustainable laboratory practices. Despite the promise of LAH, debates persist regarding its long-term efficacy compared to synthetic stabilizers. Ongoing research is needed to further elucidate its mechanisms of action and optimize its application across diverse environmental conditions [1].

## 2. Materials and Methods

### Enzyme Preparation and Preservation

Pfu DNA polymerase (1 unit/µl) was obtained from a locally produced enzyme at the Atomic Energy Commission of Syria (AECS), expressed from the Pobl6 plasmid, a gift from the Open Bioeconomy Lab at the University of Cambridge, which produces an open-source Pfu polymerase derivative [3]. Aliquots of 20 µl Pfu polymerase were prepared with or without 1 M LAH (USB Corporation). Control aliquots containing only Pfu polymerase, a 1:1 mixture of Pfu and Taq polymerase (GeneDireX), and Taq polymerase alone were also prepared. All aliquots were stored at 37°C, 25°C, and 2–8°C for three months, with control aliquots stored at -20°C to serve as a reference for optimal activity. Storage conditions are summarized in Table 1.

**Table 1:** Summary of storage temperatures, durations, polymerases, and preservatives tested.

| Temperature | Taq + Pfu | Pfu | Taq | Storage Duration |
| --- | --- | --- | --- | --- |
| 37°C | With LAH / No LAH | With LAH / No LAH | With LAH / No LAH | 3 months |
| 25°C | With LAH / No LAH | With LAH / No LAH | With LAH / No LAH | 3 months |
| 2–8°C | With LAH / No LAH | With LAH / No LAH | With LAH / No LAH | 3 months |
| -20°C | With LAH | With LAH | With LAH | 3 months |

**LAH: L-arginine hydrochloride**

### LAH: L-arginine hydrochloride

#### Enzyme Activity Assay

PCR amplification of a 1250 bp DNA fragment from the Pobl6 plasmid (Open Bioeconomy Lab, University of Cambridge) was performed using 1 µl of polymerase from each storage condition. Standard PCR conditions were applied, with the reaction mix, thermal cycling program, and primer pair designed using the Benchling platform [6], as detailed in Tables 2, 3, and 4.

**Table 2:** PCR mix components for amplifying DNA templates.

| Component | Volume (μl) |
| --- | --- |
| dNTPs | 1.5 |
| PCR buffer 10x | 2.5 |
| MgCl<sub>2</sub> | 1.5 |
| Primer F | 1.0 |
| Primer R | 1.0 |
| DNA template | 1.0 |
| DMSO | 1.0 |
| DNA polymerase | 1.0 |
| dH<sub>2</sub>O | 14.5 |
| Total Volume | 25.0 |

**Table 3:** PCR program for amplifying the 1250 bp amplicon.

| Stage | Temperature | Time | Cycles |
| --- | --- | --- | --- |
| Initial Denaturation | 95°C | 5 min | 1 |
| Denaturation | 95°C | 1 min | 35 |
| Annealing | 58°C | 40 sec |  |
| Extension | 72°C | 1 min |  |
| Final Extension | 72°C | 7 min | 1 |
| Storage | 4°C | ∞ | 1 |

**Table 4:**
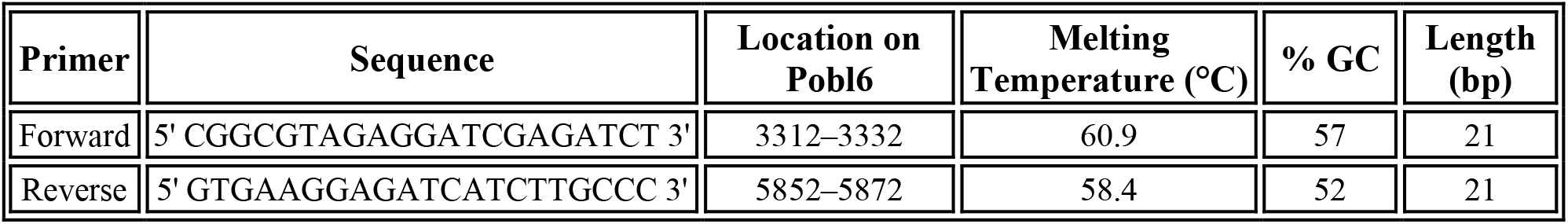
Primer pairs used in the PCR.

#### Gel Electrophoresis

PCR products were separated by 1% agarose gel electrophoresis and visualized using ethidium bromide staining, with a 1 Kb DNA ladder (Vivantis) as a marker. Gel images were captured using a UVP GelDocIT system. Relative enzyme activity was determined by comparing the presence and intensity of PCR product bands from enzymes stored at 4°C, 25°C, and 37°C, with and without LAH, to those from the -20°C control using ImageJ software for semi-quantitative analysis.

## 3. Results

### 3.1. Three-Month Shelf-Life Test Without L-Arginine Hydrochloride

To evaluate enzyme stability in the absence of LAH, PCR amplification was performed using enzymes stored without a preservative. Data were obtained from a single experiment, with consistent results across samples, though replicate experiments are needed for statistical validation. At 4°C, Pfu polymerase (sample 1), the Pfu + Taq mixture (sample 2), and Taq polymerase alone (sample 3) retained enzymatic activity, producing visible amplification bands. At 25°C, Pfu polymerase (sample 4) and the Pfu + Taq mixture (sample 5) retained activity, but Taq polymerase alone (sample 6) showed no amplification. At 37°C, none of the enzymes—Pfu polymerase (sample 7), Pfu + Taq mixture (sample 8), or Taq polymerase alone (sample 9)—produced amplification bands, indicating complete degradation. These results, shown in Figure 1, confirm that higher temperatures accelerate enzyme degradation without a stabilizer.

**Figure 1:**
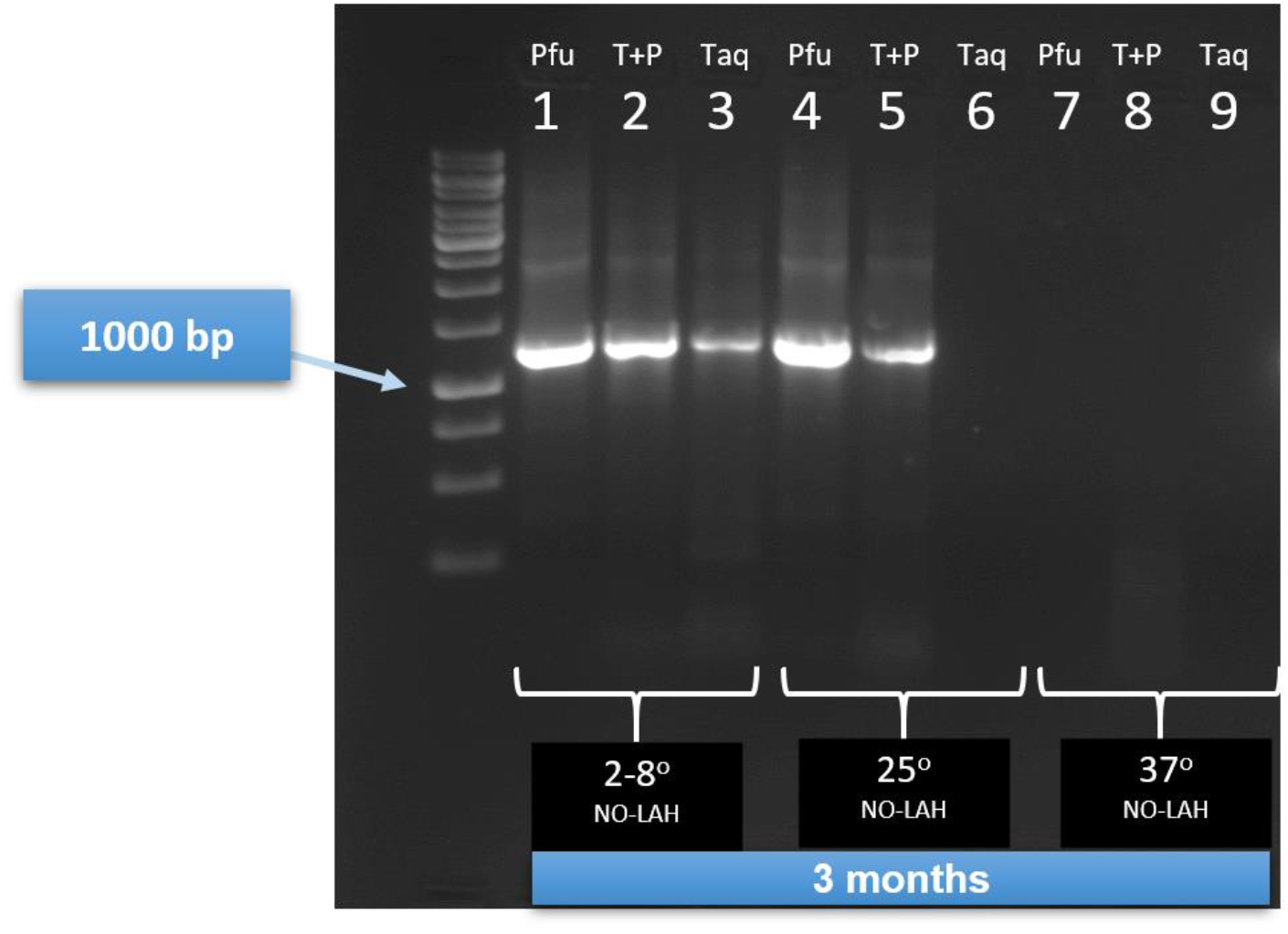
An agarose gel (1%) showing PCR amplification results for enzymes stored without L-arginine hydrochloride (LAH) for 3 months at 4°C, 25°C, and 37°C. The gel includes samples 1–3 (4°C: Pfu, Pfu + Taq, Taq), samples 4–6 (25°C: Pfu, Pfu + Taq, Taq), and samples 7–9 (37°C: Pfu, Pfu + Taq, Taq). Bands represent successful amplification of a 1250 bp fragment, with a 1 Kb DNA ladder as reference. The absence of bands at 37°C indicates enzyme degradation.

### 3.2. Three-Month Shelf-Life Test with L-Arginine Hydrochloride

Enzyme stability was assessed after three months of storage with 1 M LAH, followed by PCR amplification of the 1250 bp target fragment. Data were obtained from a single experiment, with consistent results across samples. At 4°C, Pfu polymerase (sample 10), the Pfu + Taq mixture (sample 11), and Taq polymerase alone (sample 12) produced successful amplification bands. At 25°C, Pfu polymerase (sample 13), the Pfu + Taq mixture (sample 14), and Taq polymerase alone (sample 15) also showed positive amplification. Notably, at 37°C, Pfu polymerase (sample 16), the Pfu + Taq mixture (sample 17), and Taq polymerase alone (sample 18) maintained activity, producing clear amplification bands. These results, depicted in Figure 2, demonstrate LAH’s effectiveness in preserving enzyme function across a range of temperatures.

**Figure 2:**
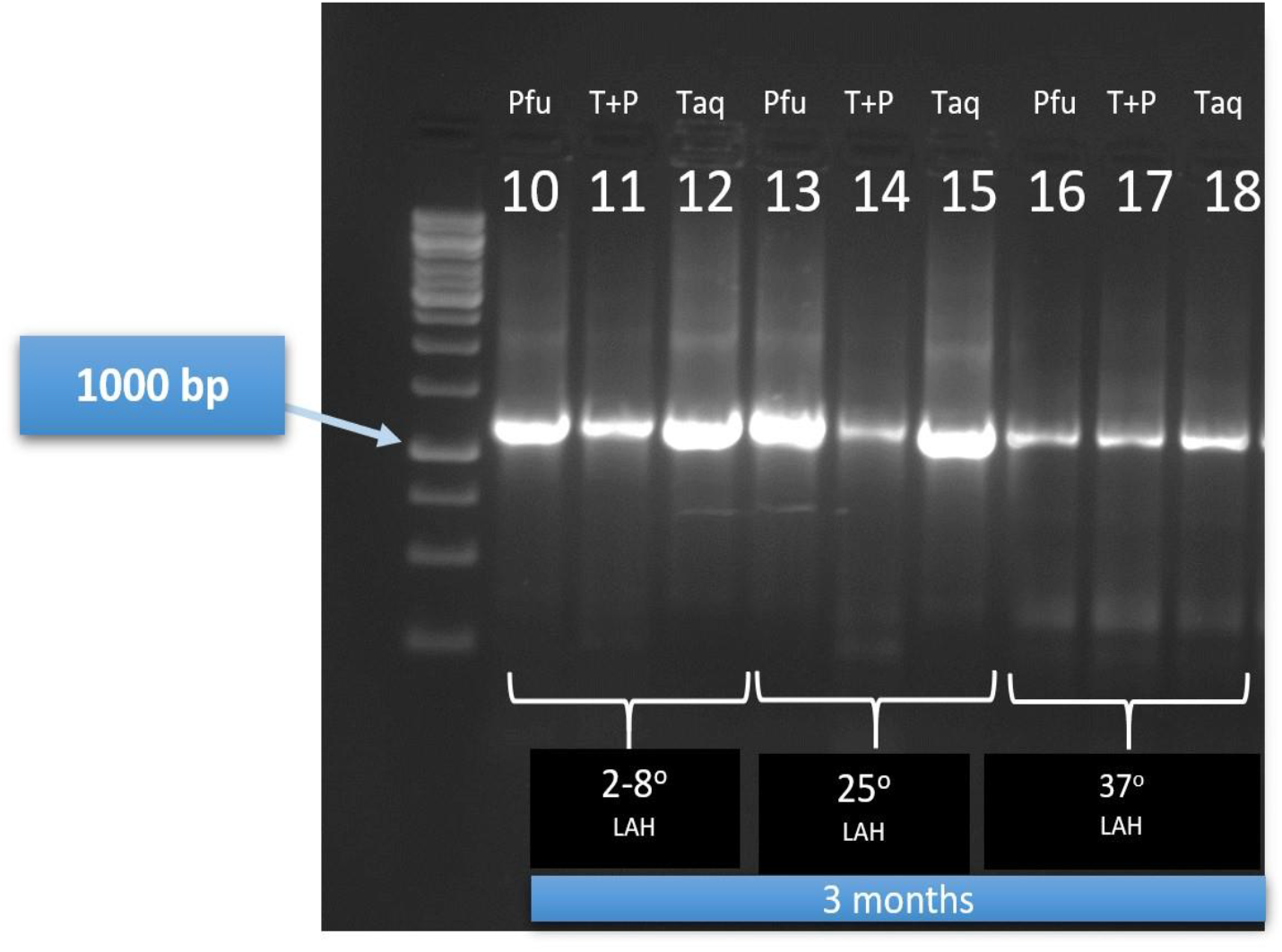
An agarose gel (1%) showing PCR amplification results for enzymes stored with 1 M L-arginine hydrochloride (LAH) for 3 months at 4°C, 25°C, and 37°C. The gel includes samples 10–12 (4°C: Pfu, Pfu + Taq, Taq), samples 13–15 (25°C: Pfu, Pfu + Taq, Taq), and samples 16–18 (37°C: Pfu, Pfu + Taq, Taq). Bands represent successful amplification of a 1250 bp fragment, with a 1 Kb DNA ladder as reference. The presence of bands at all temperatures demonstrates LAH’s stabilization effect.

### 3.3. Three-Month Viability Test with L-Arginine Hydrochloride at -20°C

Enzyme viability under freezer storage was evaluated by PCR amplification of the 1250 bp fragment using enzymes stored at -20°C. Data were obtained from a single experiment. Without LAH, Pfu polymerase (sample 22), the Pfu + Taq mixture (sample 23), and Taq polymerase alone (sample 24) retained activity. With LAH, Pfu polymerase (sample 25), the Pfu + Taq mixture (sample 26), and Taq polymerase alone (sample 27) also produced successful amplifications. A commercial Pfu enzyme (Vivantis) stored at -20°C served as a positive control (sample 21) and produced a faint amplification band. Negative controls using locally produced Pfu polymerase (-VE1, sample 19) and commercial Pfu (-VE2, sample 20) without template DNA showed no amplification, confirming the absence of contamination. These results are shown in Figure 3.

**Figure 3:**
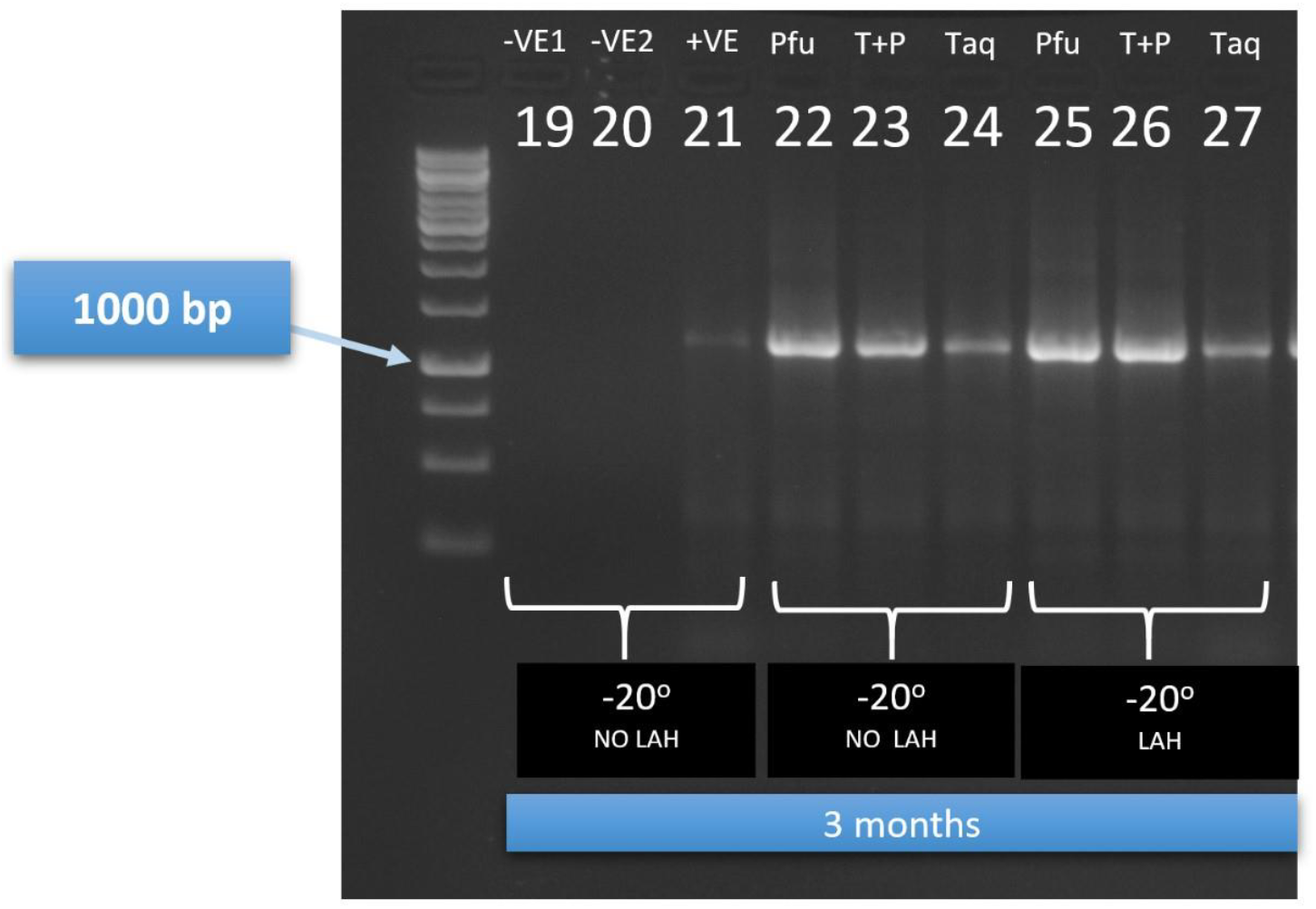
An agarose gel (1%) showing PCR amplification results for positive and negative controls and enzymes stored at -20°C with and without LAH. The gel includes samples 19–20 (negative controls: -VE1, -VE2 with no template DNA), sample 21 (positive control: commercial Pfu, -20°C), samples 22–24 (no LAH, -20°C: Pfu, Pfu + Taq, Taq), and samples 25–27 (with LAH, -20°C: Pfu, Pfu + Taq, Taq). Bands represent successful amplification of a 1250 bp fragment, with a 1 Kb DNA ladder as reference.

### 3.4. Quantitative Analysis of Enzyme Activity

Band intensity analysis using ImageJ revealed that LAH-treated enzymes retained 70–90% of their activity relative to -20°C controls across all temperatures. In contrast, enzymes stored without LAH at 37°C showed complete loss of activity, while Pfu polymerase retained partial activity (approximately 40%) at 25°C without LAH, and Taq polymerase showed no activity at 25°C or higher without LAH. The Pfu + Taq mixture exhibited intermediate stability, with LAH-treated samples retaining 75–85% activity at 25°C and 37°C. These results are summarized in Figure 4.

**Figure 4:**
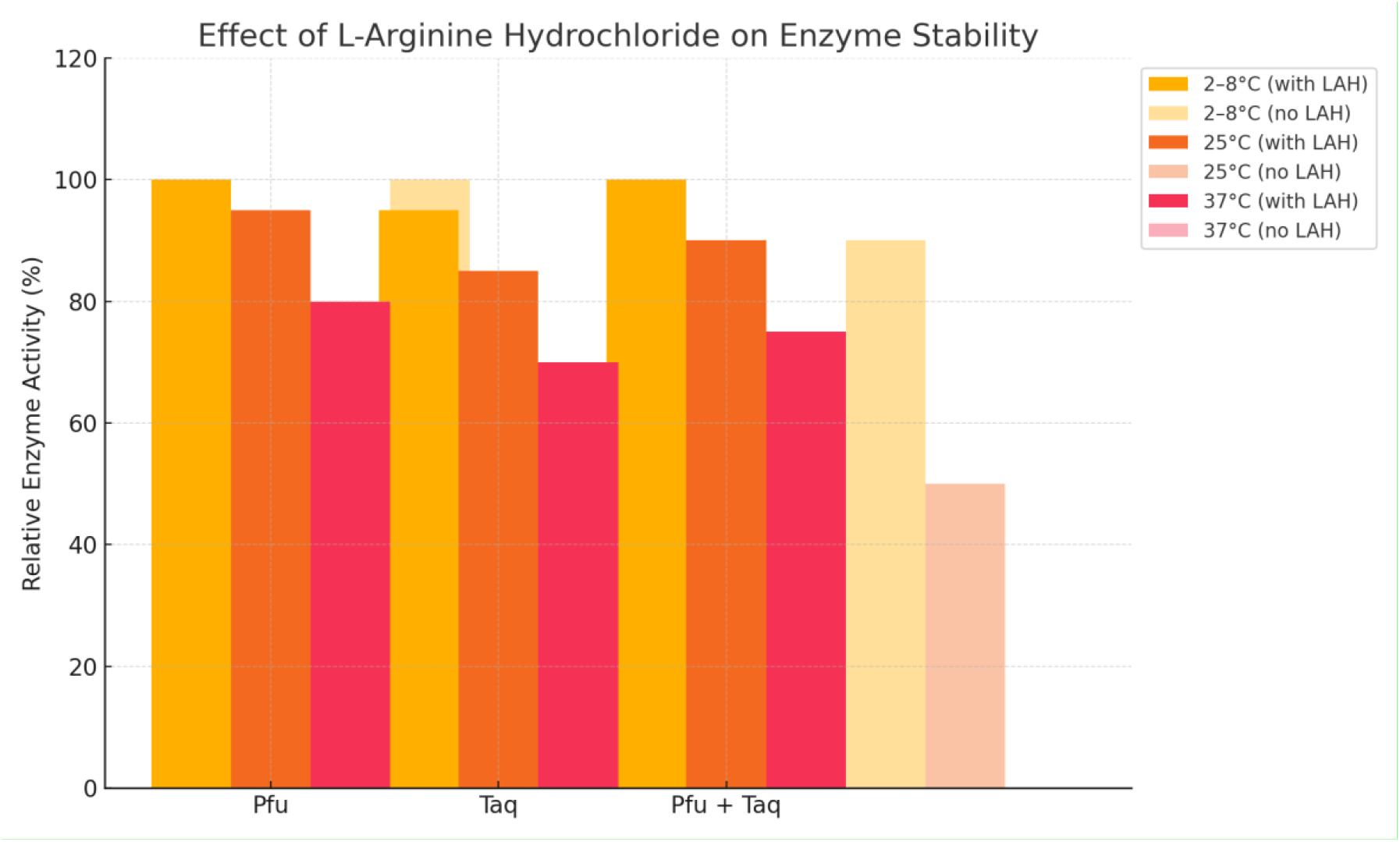
A bar chart comparing relative enzyme activity (%) after 3 months of storage with and without L-arginine hydrochloride (LAH) at 4°C, 25°C, and 37°C. Solid bars represent samples with LAH; transparent bars represent samples without LAH. Relative activity was estimated using ImageJ analysis of band intensity compared to -20°C controls. The chart highlights LAH’s preservation effect, particularly at 25°C and 37°C, with Pfu showing higher stability than Taq.

## 4. Discussion

### 4.1. Suggested Mechanisms of L-Arginine Hydrochloride in Protein Thermostability

L-arginine hydrochloride enhances protein stability through multiple molecular mechanisms, reducing aggregation, improving solubility, and maintaining structural integrity [9]. The key mechanisms include:

- **Prevention of Protein Aggregation:** LAH disrupts protein-protein interactions, preventing aggregation and ensuring active sites remain accessible, thus preserving native functional conformations. This effect is achieved by reducing hydrophobic interactions, as demonstrated by Golovanov et al. [7], ensuring that active sites remain exposed [10].
- **Enhancement of Protein Solubility:** LAH significantly improves the solubility of proteins, particularly those with low intrinsic solubility, even at low concentrations, enhancing overall stability [17].
- **Molecular Interactions and Structural Stabilization:** The guanidinium group in LAH interacts with hydrophobic protein regions, shielding them and reducing the energetic cost of exposure, thereby stabilizing protein structure [15].
- **Thermal Stabilization:** LAH increases the thermal stability of proteins, as demonstrated by elevated assembly temperatures in proteins like insulin, though its effects on thermostable enzymes like Pfu polymerase require further exploration [12].
- **Competitive Water Binding and Hydrolysis Resistance:** LAH competes with water molecules for protein interactions, increasing the energy barrier for hydrolytic degradation and enhancing stability [16].
- **Electrostatic Interactions and Charge Stabilization:** The positively charged amino groups in LAH form electrostatic bonds with negatively charged protein regions, reinforcing structural integrity and minimizing conformational changes [3].

These mechanisms collectively underscore LAH’s multifaceted role as a stabilizing agent, particularly for enzymes under thermal stress. While LAH offers a simple and effective stabilization method, other approaches, such as enzyme immobilization, have also been explored to enhance thermostability [12]. Further studies are needed to elucidate LAH’s specific effects on thermostable DNA polymerases like Pfu under diverse storage conditions.

### 4.2. Effect of L-Arginine Hydrochloride on Enzyme Stability

The results unequivocally demonstrate that LAH significantly enhances the thermal stability of Pfu polymerase, enabling it to retain activity at 37°C for three months, whereas activity was completely lost without LAH at this temperature. LAH-treated Pfu polymerase maintained 80– 90% activity at 25°C and 37°C, suggesting robust protection against thermal degradation, consistent with findings that LAH prevents protein aggregation [8]. The enhanced thermal stability observed with LAH aligns with recent findings that arginine derivatives protect enzymes under stress conditions by stabilizing their native conformation [14]. Pfu polymerase exhibited the highest intrinsic stability, followed by the Pfu + Taq mixture, which retained 75–85% activity with LAH at elevated temperatures. Taq polymerase alone was markedly less stable, retaining only 70– 75% activity at 25°C and 37°C with LAH, indicating that LAH’s efficacy varies by enzyme type and intrinsic thermal resistance.

## 5. Conclusions

This study establishes LAH as a highly effective stabilizing agent for DNA polymerases, enabling storage at ambient and elevated temperatures (up to 37°C) without significant loss of activity. These findings have profound implications for low-resource settings, where cold storage is often unavailable, by reducing cold chain dependency, enabling on-site PCR diagnostics, and lowering costs associated with enzyme importation and storage. These results align with recent advancements in enzyme stabilization for diagnostics in low-resource settings [19]. LAH offers a sustainable, cost-effective alternative to traditional preservation methods, with potential applications in global health and biotechnology.

## 6. Limitations & Future Research

While LAH demonstrates significant promise, limitations include the lack of quantitative PCR (qPCR) data, reliance on a single experiment without replicates, and a three-month study duration. Future research should focus on optimizing LAH concentrations, assessing stability beyond three months, testing additional polymerases, conducting qPCR for precise quantification, performing replicate experiments for statistical robustness, and investigating LAH’s mechanistic role at the molecular level.

## 7. Ethical Statement: Conflict of Interest

The authors declare no conflicts of interest.

## 9. Acknowledgments

We thank Dr. Jennifer Molloy from the Open Bioeconomy Lab, University of Cambridge, for providing the Pobl6 plasmids.

We also Thank for Dr. Samer Haj Kaddour for his administrative help for this research.

